# Focused ultrasound enhances sensorimotor decision-making in monkeys

**DOI:** 10.1101/041152

**Authors:** Matthew E Downs, Tobias Teichert, Amanda Buch, Maria Eleni Karakatsani, Carlos Sierra, Shangshang Chen, Elisa E Konofagou, Vincent P Ferrera

**Author notes:** Corresponding Author: Matthew E. Downs.

## Abstract

Noninvasive brain stimulation using focused ultrasound has many potential applications as a research and clinical tool. Here, we investigated the effect of focused ultrasound (FUS) combined with systemically administered microbubbles on visual-motor decision-making behavior in monkeys. We applied FUS to the putamen in one hemisphere to open the blood-brain barrier, and then tested behavioral performance 3-4 hours later. On days when the monkeys were treated with FUS, their decisions were faster and more accurate than days without sonication. The performance improvement suggested both a shift in the decision criterion and an enhancement of the use of sensory evidence in the decision process. FUS also interacted with the effect of a low dose of haloperidol. The results suggest that a two-minute application of FUS can have a sustained impact on performance of complex cognitive tasks, and may increase the efficacy of psychoactive medications. The results lend further support to the idea that the dorsal striatum plays an integral role in evidence- and reward-based decision-making.

## Introduction

Brain stimulation is an essential tool for investigating causal brain-behavior relationships, mapping brain circuits, and treating neurological disorders. Current stimulation methods are either invasive (electrical or chemical stimulation, optogenetics), or have limited penetrability (TMS) or localizability (TDCS) (Borchers *et al* 2012, Calvo and Combra, 2006, Dubuisson and Dennis, 1977, Kobayashi and Pascual-Leone, 2003, Miller, 1965, Nitsche *et al* 2003). Focused ultrasound (FUS) is emerging as a non-invasive technology capable of penetrating the skull and meninges to deliver mechanical energy to deep brain structures. FUS with systemically administered microbubbles has been shown to open the blood-brain barrier (BBB) in various animal models, and may also directly modulate neural activity (Tung *et al* 2011, Marquet *et al* 2014, Downs *et al* 2015, Chu *et al* 2015, McDannold *et al* 2005, McDannold *et al* 2012).

Recent studies in monkeys and humans have provided evidence that FUS alone can modify perception and behavior (Bystritsky *et al* 2011, Deffieux *et al* 2013, Hameroff *et al* 2013, Legon *et al* 2014, Lee *et al* 2016). Deffieux *et al* found that FUS can increase the latency of antisaccades in monkeys. Tactile discrimination was enhanced during FUS stimulation of the somatosensory cortex in human subjects, while overall mood improved when the frontal-temporal cortex was stimulated with FUS (Hameroff *et al* 2013, Legon *et al* 2014). Lee *et al* (2016) were able to evoke visual phosphenes and concomitant EEG activity. Further investigation using different species, brain targets, behavioral tasks, and FUS methodologies is warranted to establish the effectiveness and range of applications for this approach. Here, we consider whether FUS with microbubbles has an effect on the performance of a complex cognitive task 3-4 hours after treatment.

FUS with microbubbles can increase the permeability of the BBB, which remains open for up to 48 hours after treatment (Marquet *et al* 2014), raising the possibility that cognitive or behavioral changes might occur during this time period. In the current study, FUS was applied to the putamen, a part of the basal ganglia involved in cognition, reward, and movement control. We sought to devise a behavioral paradigm that would be sensitive to changes in perception, motor performance, decision-making and motivation. We therefore trained monkeys to perform a perceptual decision-making task using a touchpanel display. Monkeys learned to perform the task with either hand, and had to alternate hands during the experiment. The task involved the detection of coherent visual motion (Lappin and Bell, 1976) and also included a reward manipulation to test motivation. The advantage of this task is that it allows quantitative measures of response time and accuracy for effectors both ipsi- and contralateral to the FUS intervention. Such measures have been used to develop sophisticated computational models of decision-making in several species including humans and monkeys (Stone, 1960; Ratcliff, 1978). Electrophysiological studies point to a critical role of the striatum (caudate and putamen) in similar tasks (Ding and Gold, 2013).

We used the coherent motion detection task to investigate the effect of FUS on decision-making and motor performance. Rhesus monkeys were treated with FUS and intravenous microbubbles to open the BBB and then tested behaviorally 3-4 hours later. The current study also investigated the interaction of FUS with a low dose of the D2 dopamine antagonist haloperidol, as this technique could be used to non-invasively facilitate drug effects while minimizing side effects, or to deliver drugs that cannot cross the intact BBB.

On days when the monkeys received the FUS with microbubbles treatment, their decisions were faster and more accurate compared to days without sonication. Overall, response times were faster for the hand contralateral to the sonicated hemisphere than for the ipsilateral hand. A threshold dose of haloperidol alone reduced response time, but impaired accuracy. FUS with microbubbles enhanced the accuracy reduction and partially reversed the response time reduction. These results indicate that FUS with microbubbles can be used alone or in combination with psychoactive drugs to modify performance on complex tasks.

## Methods

All procedures with monkeys were approved by the Institutional Animal Care and Use Committees (IACUC) of Columbia University and the New York State Psychiatric Institute (NYSPI). Two adult male Macaca mulatta (N, O) were used in all experiments (9 and 20 years old, 5.5 and 9.5 kg). The monkeys were surgically naïve and underwent no procedures during the course of these experiments other than those described below. Monkeys were provided daily rations of vitamin enriched dry primate biscuits, as well as enrichment toys and allowed access to play modules. Monkeys were trained using operant conditioning to perform a visual-motor decision-making task using a touchpanel display. Prior to data collection, monkeys were trained for several months until they reached asymptotic performance. On behavioral testing days, monkeys performed the task for fluid reward until satiated. After behavioral testing, Monkeys were given a fruit treat (banana, apple, or orange). On days when behavioral testing was not conducted, monkeys were given a liter of water.

### Focused Ultrasound and Drug Delivery

On selected days, monkeys received a FUS with microbubble treatment 3-4 hours prior to behavioral testing. For the FUS procedures, subjects were sedated with ketamine (10 mg/kg) and atropine (0.04 mg/kg) and placed into a stereotaxic positioning frame under general anesthesia (isoflurane 1-2%) to ensure accurate targeting. Microbubbles (4-5 um, in-house prepared) were administered intravenously at the onset of the FUS application (single element transducer, 500 kHz, 400 kPa, 10 ms pulse length, 120 second duration; H-107, Sonic Concepts, WA, USA) (Feshitan *et al* 2009). The putamen region of the basal ganglia was targeted for all experiments. Throughout the procedure, vital signs were continuously monitored (heart rate, SPO2, mean arterial pressure, respiratory rate and end tidal CO2). After the FUS procedure there was a 3 to 4 hour recovery period allowing the monkeys to fully recover from anesthesia. After the recovery period they showed normal alertness, appetite and mobility as evidenced by their ability to walk, climb and consume food.

Haloperidol, a D2 dopamine receptor antagonist (R&D Systems, Inc., Minneapolis, MN), was used during some sessions to augment neuromodulation. Haloperidol powder was dissolved in saline and titrated to the concentration of 0.01mg/kg. On selected days, before the task began, monkeys were administered either saline or haloperidol (0.01mg/kg) intramuscularly. The injection was given 5 minutes prior to the start of behavioral testing. The threshold dose of haloperidol was determined as the maximum dose that had a minimal effect on behavioral results when the BBB was intact. The timing of events during the FUS procedure, recovery, drug injection and behavioral testing is shown in figure 1A.

**Figure 1.**
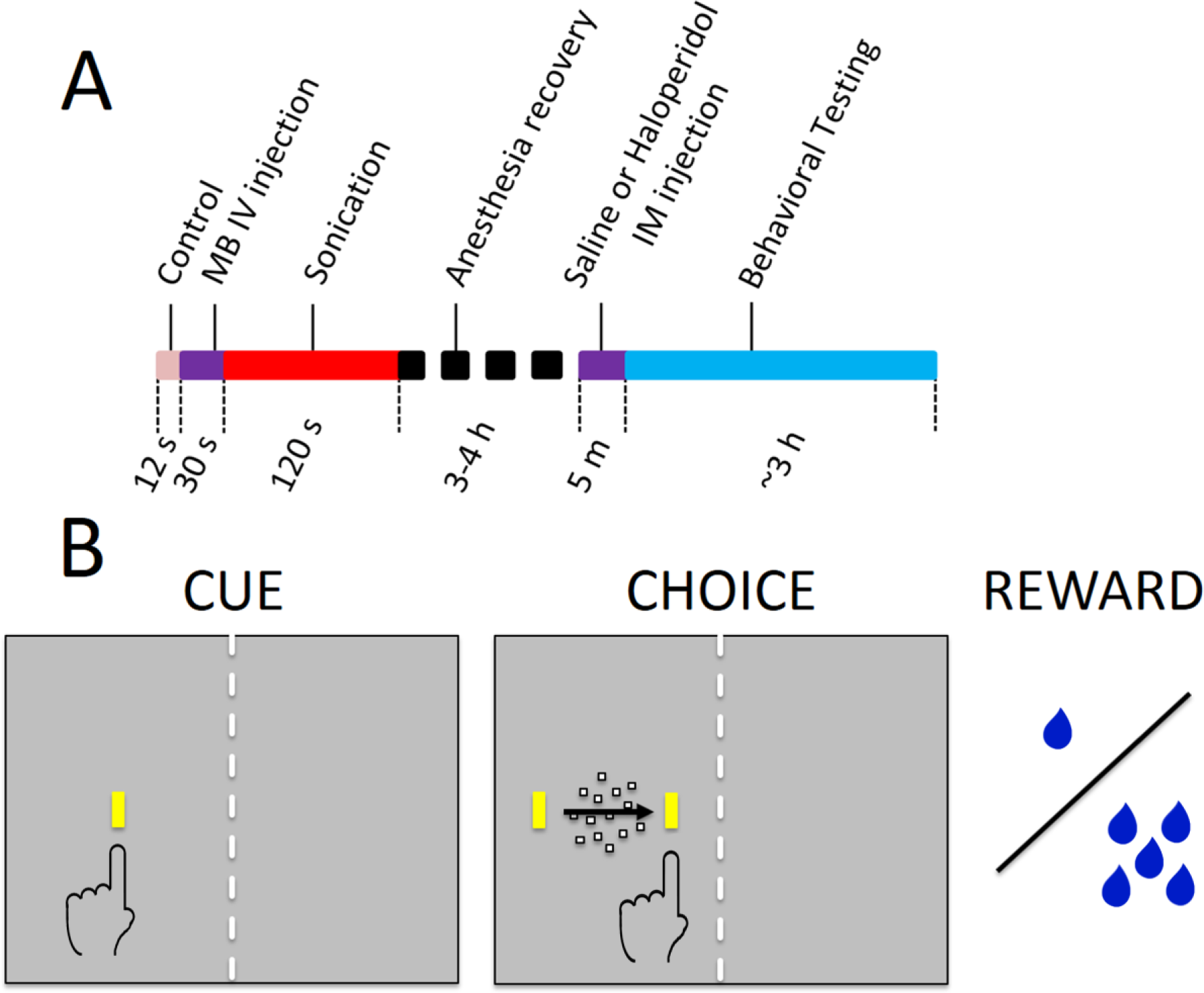
Experimental timeline and behavioral task. A) Timeline of sonication and behavioral testing. B) Decision task sequence. The monkey initiated a trial by touching the cue. A random dot motion stimulus appeared moving to the left or right, flanked by two targets. The monkey touched the target toward which the dots were moving to receive a reward. Stimuli were displayed on the right or left of the screen. A physical barrier (dashed line) forced the monkey to respond with the corresponding hand. Only the yellow bars and dot stimulus were visible to the monkey, not the blue arrow, dashed line, or hand symbol, which are used here to indicate the motion of the dots, the physical barrier separating the two halves of the screen, and the manual response, respectively. The orientation of the cue and targets indicated the size of reward (1 or 5 drops of water.)

### MRI Analysis

One day after the FUS procedure, BBB opening and safety was verified with contrast enhanced T1-weighted as well as T2-weighted MRI and susceptibility-weighted imaging scans respectively. All MRI scans (3T, Philips Medical Systems, MA, USA) were acquired 36 hours after the FUS procedure. T2-weighted (TR = 10ms, TE = 27ms, flip angle = 90°, spatial resolution = 400 x 400 μm^2^, slice thickness = 2 mm with no interslice gap) and susceptibility-weighted image (TR = 19ms, TE = 27ms, flip angle = 15°, spatial resolution = 400 x 400 μm^2^, slice thickness = 1 mm with no interslice gap) scans were used to verify the safety of the procedure. Contrast enhanced T1-weighted (TR = 19ms, TE = 27ms, flip angle = 15°, spatial resolution = 400 x 400 μm^2^, slice thickness = 1 mm with no interslice gap) scans were acquired 30 minutes after IV administration of 0.2ml/kg gadodiamide (Omniscan®, 573.66 DA, GE, Healthcare, Princeton, NY, USA). Gadodiamide was used as the contrast agent as it does not cross the intact BBB. All acquired scans were aligned with a previously acquired stereotactically aligned structural T1-weighted MRI scan to verify opening in the targeted region. The contrast enhanced T1-weighted scans were then post processed to quantify the volume of opening. This process has been thoroughly discussed elsewhere (Downs *et al* 2015).

### Behavioral Testing

Monkeys sat in a custom-made polycarbonate primate chair that allowed them to reach out to visual stimuli presented on a 20-inch LCD touchscreen monitor (NEC 2010x with 3M SC4 resistive touchscreen) placed directly in front of the chair. The resolution of the LCD was 1280 horizontal x 1024 vertical pixels (55.4 x 45.4 deg. visual angle at 14 in viewing distance) with a refresh rate of 60 Hz. The touchscreen device had a resolution of 1024 x 1024 pixels and a sampling rate of 60 Hz. The primate chair incorporated a polycarbonate midline divider so that stimuli presented on the right side of the display could only be reached by the right hand, and likewise for the left side. Behavior was reinforced with drops of fluid delivered by a juice tube mounted on the chair.

The behavioral task was presented as discrete trials lasting roughly 5 seconds each. Each trial began with a visual cue stimulus presented on the left or right side of the monitor (Figure 1B, “Cue”). The cue was a vertically or horizontally oriented yellow bar (1x3 deg, 43.8 cd/m2 luminance). The monkey touched the cue with the corresponding hand to initiate the trial. After a short delay, the cue was replaced by a random dot motion stimulus (Figure 1B, “Choice.”) The motion stimulus consisted of 100 dots (each dot was 0.17 deg square, luminance 71.6 cd/m2) moving within a circular aperture of 10 deg diameter. Some of the dots moved in random directions while others moved coherently in a single direction (dot lifetime was 2 frames). The coherent direction, either leftward or rightward, varied from trial to trial. The strength of the motion stimulus (aka motion coherence) varied from 0 to 0.7 in steps of 0.1. A particular coherence level was selected randomly for each trial and the coherence was constant for the duration of the trial. The motion stimulus was flanked on either side by two target stimuli that appeared simultaneously with the motion stimulus. The target stimuli were yellow bars that had the same orientation, size and luminance as the cue. The direction of the coherent dots indicated which target would be rewarded. The monkey was reinforced with drops of water for touching the appropriate target (Figure 1B, “Reward.”) There was no punishment for incorrect responses or failures to respond. No signal instructed the monkeys when to respond; rather, they were allowed to touch at any time after the motion stimulus and targets appeared.

To test motivation, the experiment included two reward sizes, one of which was chosen randomly on each trial: small offered reward (1 drop of water, 0.03 ml) and large offered reward (5 drops, 0.15 ml). Offered reward level on each trial was signaled by the orientation of the cue and target stimuli. Horizontal orientation indicated large reward, vertical indicated small reward.

One seventh of the trials were controls that were identical to the other trials except that the target for the incorrect response was not presented. On these trials, the monkey could ignore the motion stimulus and simply touch the correct target to receive a reward. The purpose of these trials was to assess movement accuracy and response time when no decision was required.

The complete task design thus had the following variables: display side (left or right), cue/target orientation (vertical or horizontal, corresponding to small and large reward), motion direction (left or right), motion coherence (0.0 to 0.7), and number of targets (1 or 2). This resulted in a balanced design comprising 48 conditions per block of trials. All conditions were randomly interleaved within each behavioral session.

### Statistics

Quantitative analyses were performed using Matlab 8.3 with the Statistics 9.0 toolbox (Mathworks, Natick MA.) Response times were analyzed with multivariate ANOVA and generalized linear model regression (using the glmfit function in the Matlab Statistics toolbox.) The GLM model equation was:

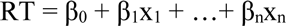

where the x_i_ are the explanatory variables described below and the β_i_ are the regression coefficients.

Performance accuracy or outcome (correct, incorrect) was analyzed with multivariate ANOVA and logistic regression (using the mnrfit function in the Matlab Statistics toolbox.) The logistic regression equation was:

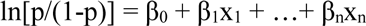

where p is the probability of a correct outcome, the xi are the explanatory variables, and the ×i are the regression coefficients. The explanatory variables used in all analyses were: subject (N, O), motion coherence (0 to 0.7, 8 levels), offered reward (1 or 5 drops), presence of sonication, sonicated hemisphere (ispsilateral or contralateral to responding hand), and drug treatment (saline or haloperidol).

## Results

### Effects of FUS on blood-brain barrier

The BBB was targeted in the putamen region of the basal ganglia for all FUS procedures. In figure 2, the red/yellow areas specify where the contrast agent was able to pass into the parenchyma, indicating successful BBB opening. The blue shaded regions indicate the region targeted by the FUS transducer. All openings achieved within this study fell inside the targeted region and no untargeted BBB openings were observed. No damage from the FUS procedure was detected; T2-weighted MRI and susceptibility-weighted imaging scans were used to detect edema but did not display any hyper- or hypointense voxels in the targeted regions.

**Figure 2.**
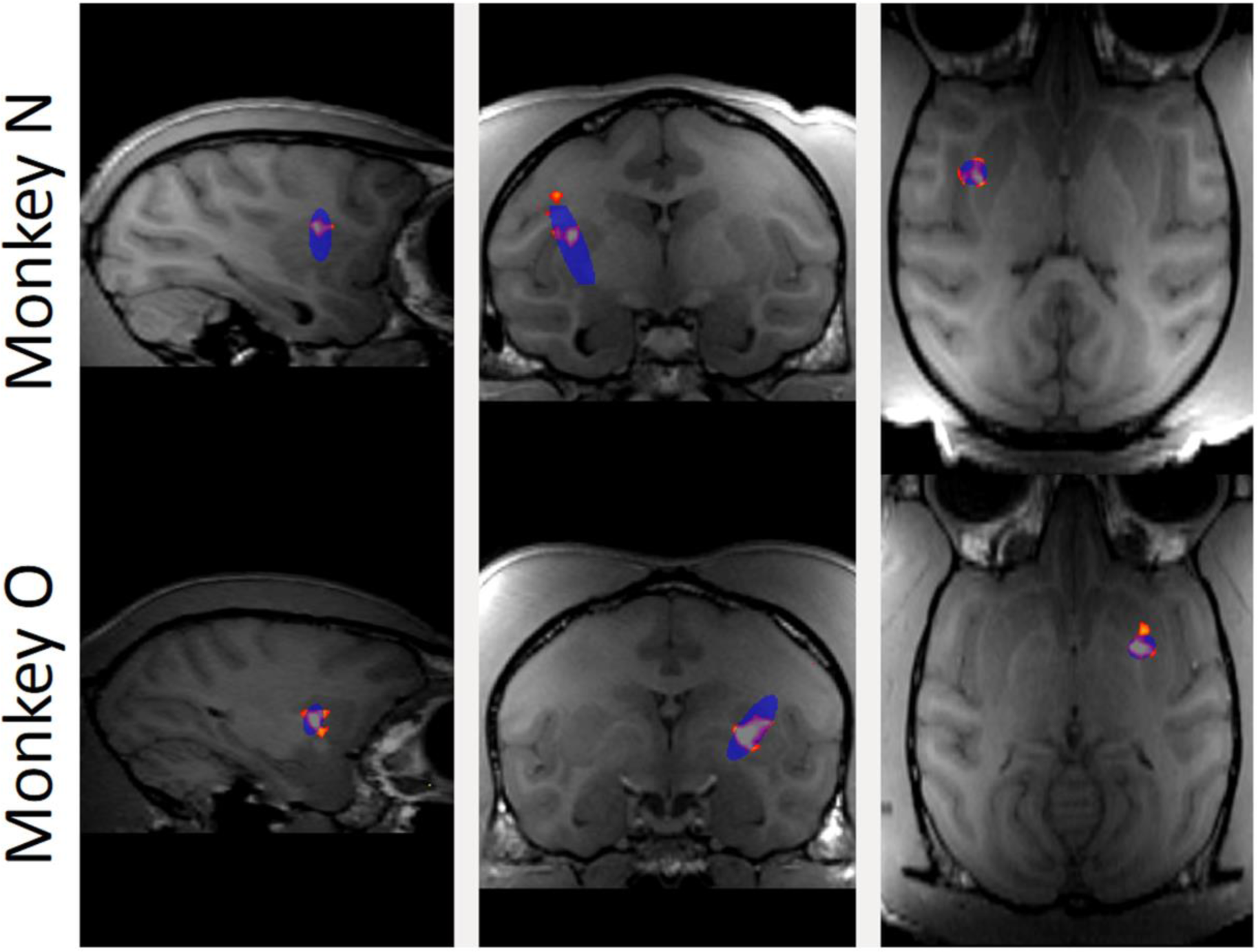
Contrast enhanced (gadodiamide) MRI of BBB opening in putamen. Top row shows sagittal, coronal and horizontal slices through the brain of monkey N. Blue oval indicates the planned target region. Red and orange voxels indicate actual BBB opening. Bottom row shows the same for monkey O.

### Effects of FUS on decision-making performance

Reward and decision uncertainty are thought to engage the dorsal striatum (Ding and Gold, 2013, Feng *et al* 2009, Lauwereyns *et al* 2002, Hikosaka *et al* 2006, Hikosaka, 2007). Thus, the behavioral task was designed to test the ability of monkeys to make decisions based on uncertain sensory evidence and variable rewards. Two monkeys performed the motion detection task during a total of 31 behavioral sessions (16 for monkey N, 15 for monkey O.) N completed an average of 1385 trials per session (22,154 total trials), while O averaged 931 trials (13,960 total).

Behavior was quantified in terms of response time, touch accuracy, and decision accuracy. Response time was measured as the interval between motion stimulus/target onset and the first touch. Touch accuracy was the distance from the center of the target to the point of first contact registered by the touchpanel. Decision accuracy was measured as the percent correct choices relative to total correct and incorrect responses. Results for the two monkeys were qualitatively similar, except that monkey N (the younger of the pair) tended to respond faster and more accurately overall.

Each trial began with the presentation of a cue stimulus (Figure 1B), which the monkey could touch to proceed with the trial. The response time and spatial topography of this initial touch provide an indication of whether any of the experimental manipulations resulted in a simple motor deficit. Touch error was quantified as the spatial dispersion of the initial touches about their mean as well as the radial distance from the center of the cue to the location of the first touch. Figure 3A shows the two cue locations and the initial touch locations, separated by sonication condition. The centroid of each ellipse is the mean touch location and the size of the ellipse is proportional to the dispersion about the mean (standard deviation). Although there were small systematic differences between conditions, the touches were tightly clustered in all conditions, providing evidence that touch accuracy was unimpaired by sonication.

**Figure 3.**
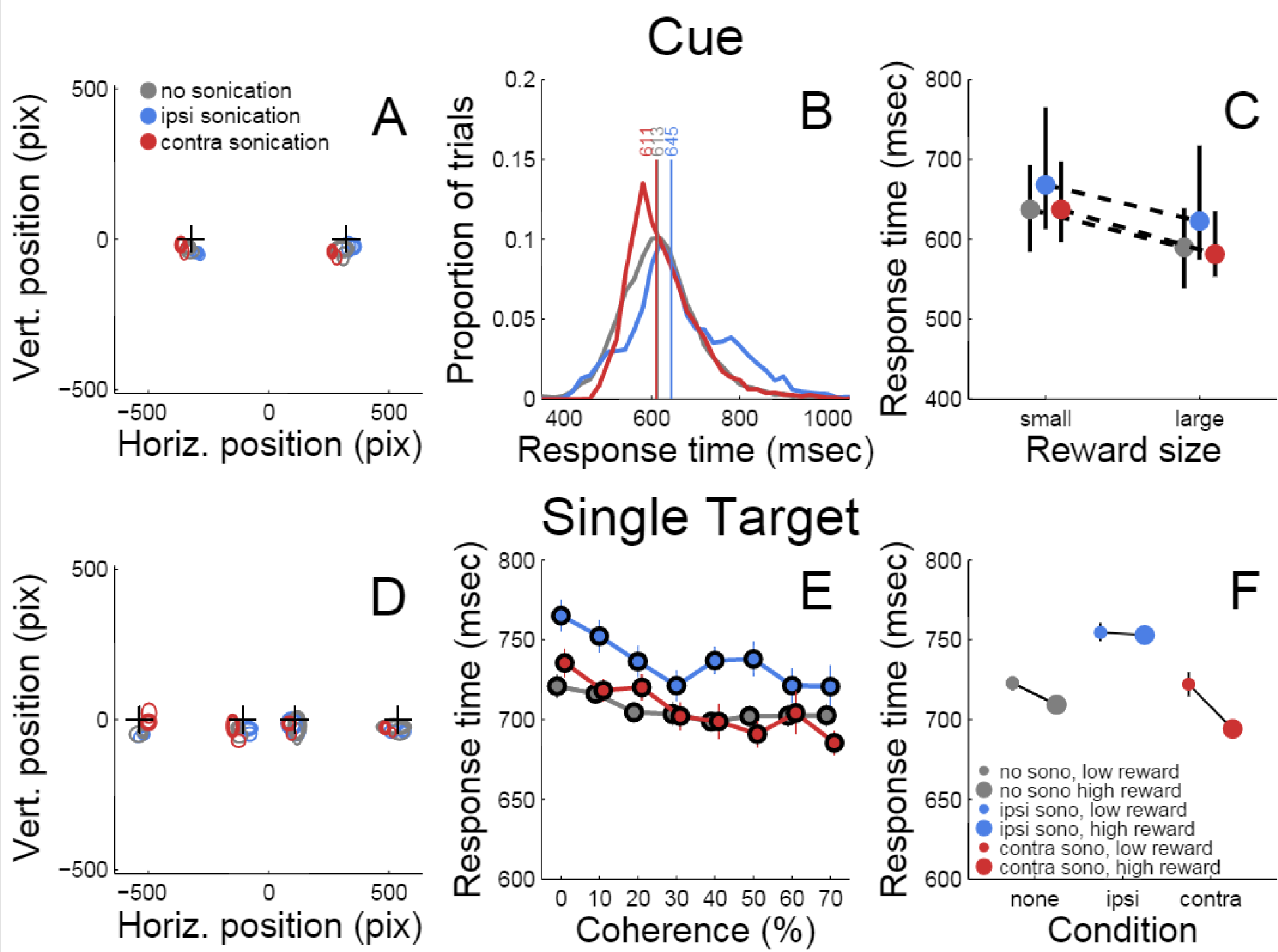
Responses to initial cue and single target. A) Accuracy of touches to initial cue. Plusses (+) indicates cue positions. Ellipses indicate mean location and dispersion of initial touches sorted by condition (legend). B) Distributions of initial touch response times sorted by condition (color code as in A). Vertical lines and numbers indicate the median of each distribution. Color code is same as panel A. C) Dependence of response time on reward size and sonication. Symbols indicate median response time (color code as in A), black lines indicate interquartile range. D). Accuracy of touches on trials with single targets (same conventions as A). E) Response time to single targets as a function of motion strength (coherence) and sonication (colors as in A) F) Effect of reward and sonication on response times (mean ± s.e.).

Table 1 summarizes the effects of reward, sonication and drug on initial touch error. All of the main effects were significant. However, the magnitudes of the errors, as assessed by the GLM analysis, were generally small. For example, reward size was correlated with touch error, but the effect amounted to 10 pixels (2.9 mm), which is less than the width of the monkey’s fingertip. The effects of sonication and drug on touch error were roughly half as large (Table 1).

**Table 1.** Multivariate ANOVA and GLM analysis of motor error and response time to the cue for all sessions. Explanatory variables were subject, offered reward (rew), sonication (sono), and haloperidol (drug). Reward, sonication and haloperidol were nested within subject. * = p<0.05, ** = p<0.01, *** = p<0.001, **** = p<0.0001

| Cue Response Error (n = 36112 trials) |  |  |  |  |  | Cue Response Time (n = 36112 trials) |  |  |  |  |
| --- | --- | --- | --- | --- | --- | --- | --- | --- | --- | --- |
|  | *ANOVA* |  |  | *GLM* |  | *ANOVA* |  |  | *GLM* |  |
| *EV* | *F* | *p* | *df* | *beta* | *p* | *F* | *p* | *df* | *beta* | *p* |
| subject | 28.6 | **** | 1 | -1.8 | * | 0.83 | 0.36 | 1 | 3.0 | * |
| rew | 1192.1 | **** | 2 | -10.4 | **** | 1028.6 | **** | 2 | -52.5 | **** |
| sono | 276.0 | **** | 2 | 5.2 | **** | 349.2 | **** | 2 | 32.5 | **** |
| drug | 525.5 | **** | 2 | -4.1 | **** | 21.8 | **** | 2 | 6.7 | **** |
| rew x sono | 124.9 | **** | 2 |  |  | 40.4 | **** | 2 |  |  |
| rew x drug | 12.3 | **** | 2 |  |  | 19.5 | **** | 2 |  |  |
| sono x drug | 97.3 | **** | 2 |  |  | 15.6 | **** | 2 |  |  |

Response time was defined as the period between the appearance of the cue and the first contact registered by the touch panel. The response time distributions are shown in figure 3B, sorted by sonication condition. Sonication was associated with slower responses overall, but this was mainly due to slowing for the ipsilateral hand (32 ms), while responses with the contralateral hand were slightly faster on days with sonication than days without. When the cue indicated a large reward, responses were ~50 ms faster than for small rewards (Figure 3C and Table 1).

After the cue was touched, there was a short, random delay and then the motion stimulus appeared together with the response target(s). On one-seventh of the trials there was only one response target whose location was congruent with the motion direction. Therefore these trials did not require a decision. The motor error (Figure 3D) for single target touches tended to be only slightly larger than for cue touches (Figure 3A). Mean response time (Figure 3E) was affected by sonication, with the largest effect being a significant slowing for touches with the hand ipsilateral to the sonicated hemisphere. With sonication, the average response time with the contralateral hand (735 ms, n=1093,) was significantly faster than with the ipsilateral hand (772 ms, n=1103 t-test p<0.0001.) Responses were also significantly faster when a large reward was available and this effect interacted significantly with sonication (Figure 3F.) Statistical results (ANOVA and GLM) are given in Table 2.

**Table 2.** Multivariate ANOVA and GLM analysis of response time for all sessions. Explanatory variables were subject, motion coherence (coh), offered reward (rew), sonication (sono), and haloperidol (drug). Coherence, reward, sonication and haloperidol were nested within subject. * = p<0.05, ** = p<0.01, *** = p<0.001, **** = p<0.0001

| One Target RT (n = 4243 trials) |  |  |  |  |  | Two Target RT (n = 31869 trials) |  |  |  |  |
| --- | --- | --- | --- | --- | --- | --- | --- | --- | --- | --- |
|  | ANOVA |  |  | GLM |  | ANOVA |  |  | GLM |  |
| EV | F | p | df | beta | p | F | p | df | beta | p |
| subject | 0.1 | 0.79 | 1 | 0.81 | 0.81 | 0.1 | 0.79 | 1 | 1.7 | 0.41 |
| coh | 3.9 | **** | 14 | -40.9 | **** | 408.5 | **** | 14 | -280.2 | **** |
| rew | 12.0 | **** | 2 | -15.3 | **** | 944.7 | **** | 2 | 78.9 | **** |
| sono | 19.3 | **** | 2 | 15.0 | *** | 16.0 | **** | 2 | -8.5 | *** |
| drug | 9.8 | *** | 2 |  | * | 305.1 | **** | 2 | -37.2 | **** |
| coh x rew | 0.5 | 0.91 | 14 |  |  | 66.1 | **** | 14 |  |  |
| coh x sono | 0.8 | 0.69 | 14 |  |  | 2.7 | *** | 14 |  |  |
| coh x drug | 0.7 | 0.78 | 14 |  |  | 10.8 | **** | 14 |  |  |
| rew x sono | 0.03 | 0.97 | 2 |  |  | 127.6 | **** | 2 |  |  |
| rew x drug | 3.0 | 0.05 | 2 |  |  | 11.9 | **** | 2 |  |  |
| sono x drug | 12.6 | **** | 2 |  |  | 33.1 | **** | 2 |  |  |

Trials with two response targets required a decision, and, therefore, both decision accuracy (percent correct) and response time were analyzed. Decision accuracy improved with increasing motion coherence (Table 2 and Figure 4A). The psychometric function was fit with a Naka-Rushton function, which was then used to find the 75% correct detection threshold (Figure 4A). Thresholds for detecting motion direction were significantly lower on days when the monkeys received sonication (Figure 4B). Logistic regression results are shown in Table 3 and indicate that both sonication and larger expected reward size improved decision accuracy.

**Figure 4.**
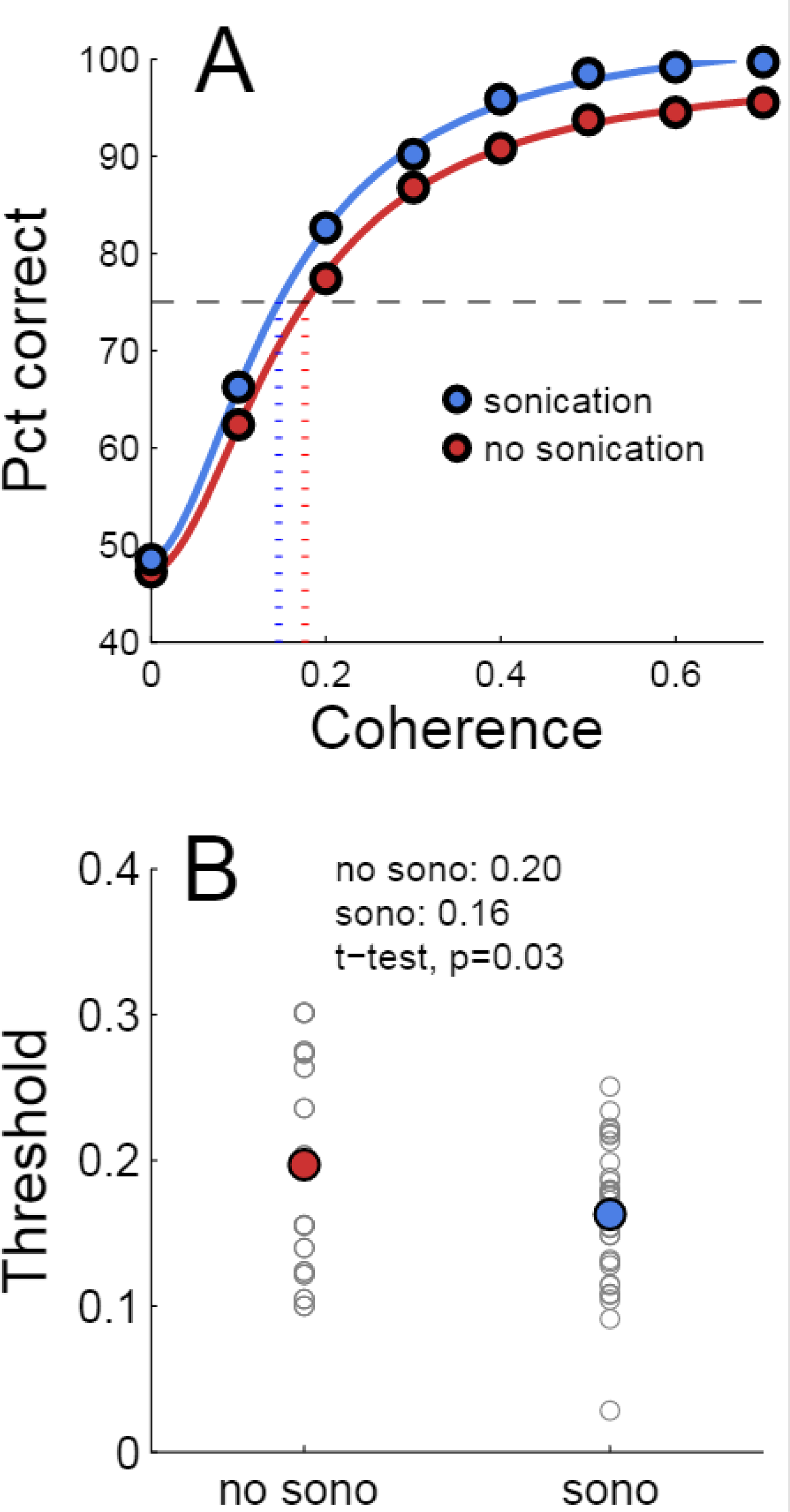
Effects of sonication on decision accuracy. A) Accuracy (percent correct) vs. coherence for sonicated and non-sonicated sessions. Solid curves are fits of Naka-Rushton functions. Dashed horizontal line indicates 75% correct level. Dotted vertical lines are coherence thresholds for 75% correct performance. B) Thresholds (75% correct) sorted by sonication condition. Small black dots are individual sessions, large colored dots are mean threshold across sessions. Note that the thresholds estimated from the aggregated data in A are not expected to precisely match the means of the individual session thresholds in B due to nonlinearities in the fitting process.

**Table 3.** Logistic regression analysis of decision accuracy. Explanatory variables were subject, motion coherence (coh), offered reward (rew), sonication (sono), and haloperidol (drug). Coherence, reward, sonication and haloperidol were nested within subject. Dependent variable was outcome (correct, incorrect). * = p<0.05, ** = p<0.01, *** = p<0.001, **** = p<0.0001

| Two Targets |  |  |  |  |  |  |
| --- | --- | --- | --- | --- | --- | --- |
|  | All sessions |  | Sessions without haloperidol |  | Sessions without sonication |  |
| *EV* | *beta* | *p* | *beta* | *p* | *beta* | *p* |
| subject | -0.60 | **** | -0.66 | **** | -0.74 | **** |
| coh | 6.24 | **** | 6.31 | **** | 5.44 | **** |
| rew | 0.26 | **** | 0.21 | **** | 0.32 | **** |
| sono | 0.17 | **** | 0.16 | **** |  |  |
| drug | -0.11 | *** |  |  | -0.11 | * |

Response times on choice trials showed a large effect of motion strength (Table 2 and Figure 5a), as found in previous studies (Roitman and Shadlen, 2002). Offered reward size had a significant effect on response time; monkeys were slower to respond when there was a larger reward at stake. This was in contrast to their behavior on single-target trials where large rewards were associated with faster response times. The results suggest that larger rewards induced the monkeys to spend more time accumulating evidence to make more accurate decisions. This is consistent with a speed-accuracy trade-off that can be accounted for by a criterion shift in sequential sampling models of decision-making (Wald, 1945).

**Figure 5.**
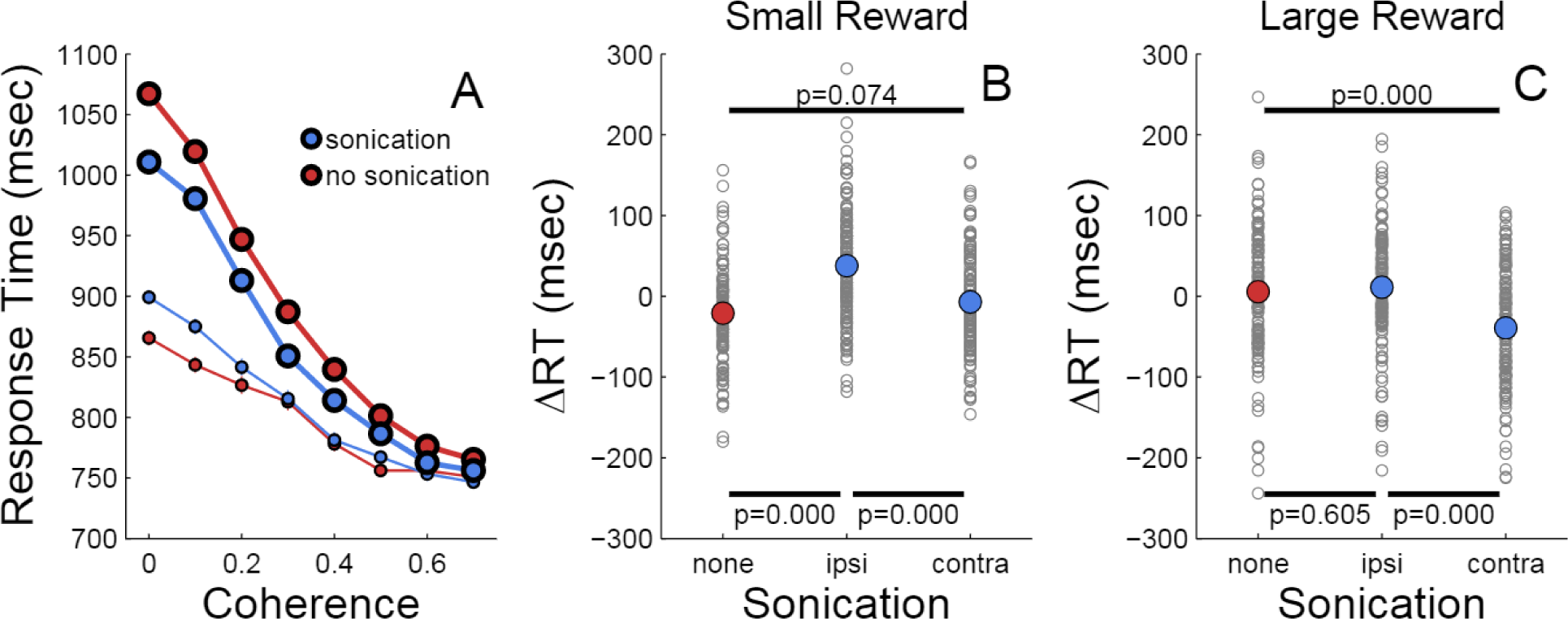
Effects of motion coherence, reward, and sonication on response time for choice (2-target) trials. A) Response time for two-target trials sorted by coherence level and sonication condition. Symbol size indicates reward size. B) Relative response time for trials sorted by session and coherence level. ΔRT is the difference between the RT segregated by condition (sono or no sono) and the overall RT. (+) and (-) on abscissa indicate presence or absence of sonication, respectively. Small symbols are for small reward trials, large symbols for large reward trials. P-values are results of t-tests and the black horizontal lines indicate the conditions that are compared. C) Relative response time for large reward trials sorted by session and coherence level.

As indicated in Table 2, the main effects of coherence and reward size on response time were highly significant. The main effect of sonication, while significant, was smaller due to a significant interaction of sonication with reward size. On small reward trials, sonication was associated with longer response times and higher accuracy. On large reward trials, sonication also improved accuracy, but reduced response times. Averaging over all conditions, the overall effect of sonication was to shorten choice response times for the contralateral hand (mean 882 ms, n=8143) compared the ipsilateral hand (929 ms, n=8160). Comparing response times for the ipsi- and contralateral hands serves as a within-session control and responses were significantly faster with the contralateral hand for both reward sizes (Figure 5 B,C.)

Haloperidol, in the absence of sonication, had significant effects on accuracy and response time compared to saline controls. For cue touches, haloperidol reduced touch error and increased response time slightly (Table 1. Error reduction: 4.2 pixels or 1.3 mm, response time increase: 6.9 ms.). Haloperidol had no significant effect on response times to single targets (Table 2.) For choice trials, haloperidol reduced both response time (Table 2) and decision accuracy (Table 3) significantly. The effects of haloperidol were thus opposite to those of reward size where larger rewards were associated with longer RT and higher accuracy, suggesting that the drug reduced motivation, consistent with the action of a D2 dopamine antagonist (Acquas *et al* 1989). Figure 6A shows decision accuracy separated by hand, reward size and coherence level (32 conditions). The haloperidol-associated reduction in accuracy was greater for sessions with sonication (red) than for those without (blue), but the difference was not significant (t-test, paired by condition, p=0.061, n=32). Figure 6B shows the distributions of the difference (haloperidol-saline) in decision accuracy for sonication (red) vs. no sonication (blue) sessions.

**Figure 6.**
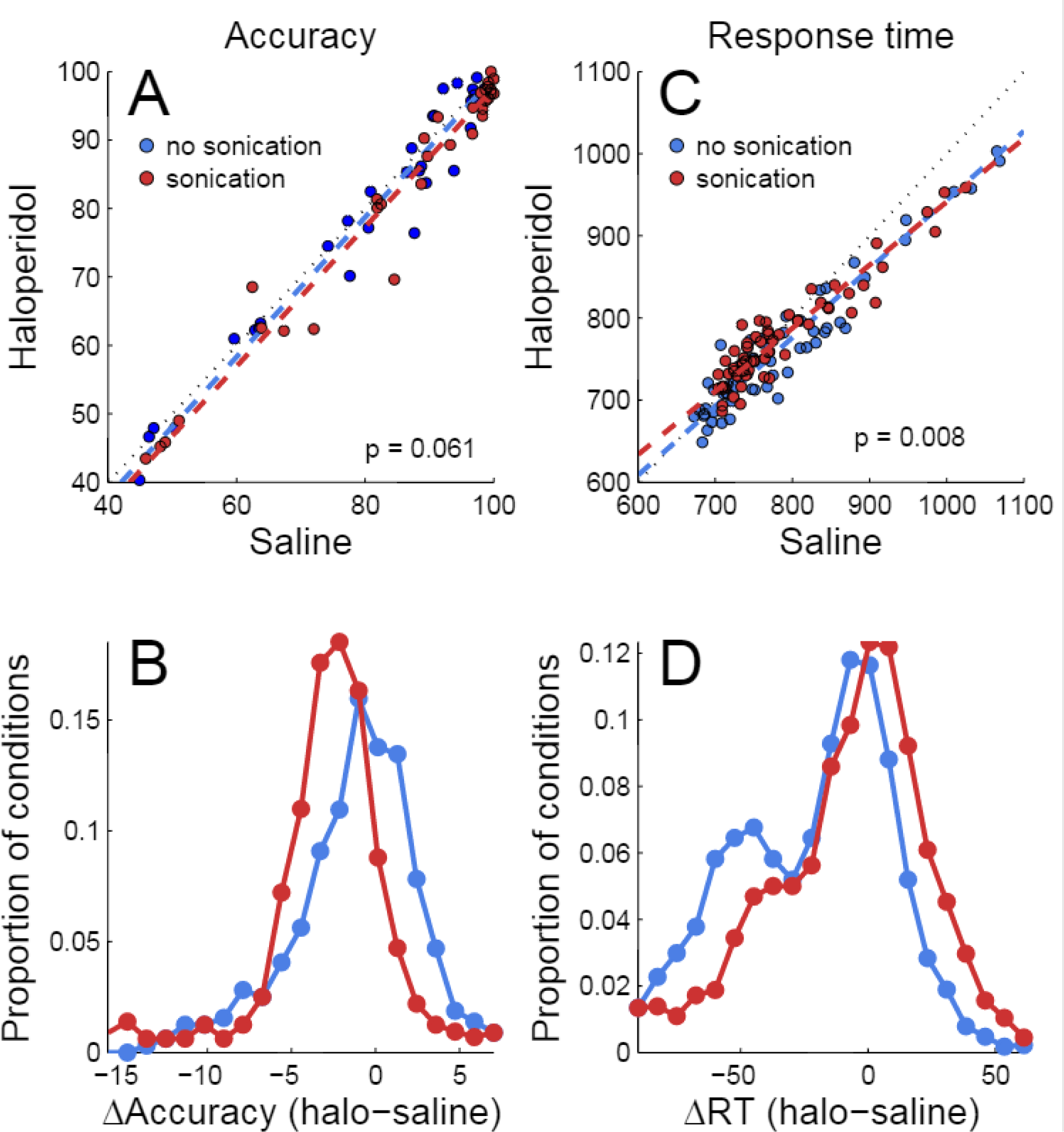
Effects of sonication and haloperidol on performance. A) Effects on decision accuracy for sessions with haloperidol vs. sessions with saline. Dashed colored lines are linear regression fits (first principal component). Black dotted line is 45 degree diagonal (x=y). “p” is the significance of a paired t-test (haloperidol vs. saline, paired by experimental condition, n = 64). B) Distributions of accuracy differences (haloperidol-saline) for sonication (red) or no sonication (blue). C) Effects on response time. Same conventions as A. D) Distributions of response time differences (same conventions as C.)

For response time, the interaction of sonication with drug was significant according to ANOVA (Table 2). Haloperidol reduced response time for both one- and two-target trials, but the reduction was smaller for sonication sessions. Figure 6C shows the effect of haloperidol compared to saline with and without sonication (data are segregated by hand, reward size, coherence level and number of targets, paired t-test p= 0.008, n=64 conditions). The distribution of the response time differences (haloperidol – saline) is plotted in figure 6D (data for both one- and two-target trials are included.)

## Discussion

We targeted the putamen in monkeys with focused ultrasound plus intravenous microbubbles to open the blood-brain barrier. Because the BBB remains open for up to 48 hours (Marquet *et al* 2014), it was important to ascertain if there are subtle cognitive or behavioral changes subsequent to the procedure.

Decision-making in monkeys has been studied previously with random dot motion tasks very similar to that used in the current study (Roitman and Shadlen 2002, Feng *et al* 2009.) Electrophysiological evidence suggests that the dorsal striatum (caudate and putamen) plays a role in such tasks (Ding and Gold 2013), thus motivating us to use a variation of the task that could reveal changes in perception, motor control, decision-making and motivation.

Sessions without sonication were used to establish a behavioral baseline. We confirmed previous work showing that response times vary inversely with the strength of the motion signal (Roitman & Shadlen 2002.). The lengthening of response times (RT) is an effective strategy to optimize accuracy, as temporal integration of weak motion signals improves decision accuracy. Manipulating the relative reward size for the two response alternatives can introduce a response bias (Feng *et al* 2009; Teichert & Ferrera 2010). Here we found that manipulating reward size for correct responses induced animals to trade response speed for accuracy, but did not introduce a response bias as there was never any incentive to choose the incorrect target. We found that when a larger reward was offered, monkeys responded significantly more slowly than they did for smaller rewards, gaining a small amount of additional accuracy by doing so. This speed-accuracy trade-off can be modeled as a criterion shift in sequential sampling models (Wald 1942, Ratcliff 2002.)

Applying FUS with microbubbles to the putamen of monkeys resulted in significant improvements in decision-making performance. Monkeys responded faster and more accurately when tested on days with sonication than on days without. Increased accuracy coupled with shorter reaction times suggests an improvement in the quality of sensory evidence or more perfect temporal integration of the motion signal. Comparison of responses with the hands ipsilateral and contralateral to the sonicated hemisphere provides a within-session control. The effects of sonication depended on the hand used to respond with response times being significantly faster for the contralateral than ipsilateral hand. Because the hand used to respond varied randomly from trial to trial, these effects are unlikely to be due to general arousal or non-specific effects of anesthesia.

Response times also depended on reward size. In the absence of sonication, choice response time was as much as 200 ms slower on large than on small reward trials, suggesting that the availability of a large reward led to more deliberative (less impulsive) decisions. The effect of reward size during choice trials was opposite to that on non-choice trials where responses were significantly faster for larger rewards.

Sonication reduced the effect of reward on choice RT by reducing RT on larger reward trials and increasing RT on small reward trials, all while increasing accuracy. Thus, for large rewards, sonication appears to improve the efficiency of decision-making, possibly by improving the quality of the sensory signal or the rate of evidence accumulation. In other words, improved decision efficiency after sonication might result from greater signal-to-noise or by reducing the leakiness of the integrator. These findings provide new evidence that the dorsal striatum (caudate and putamen) is involved in sensory evidence accumulation and thus plays an integral role in the decision process (Ding and Gold 2013.)

Decision-making performance improvements were found even though animals were tested 3-4 hours after sonication, suggesting that there may be a persistent effect on the activity or responsiveness of putamen neurons, which, in turn, may be due to a direct effect of ultrasound or an indirect effect of opening the blood-brain barrier. It is likely that BBB opening alters the local extracellular milieu, possibly by enriching the parenchymal concentration of oxygen and glucose. Ultrasound may also directly affect the permeability of mechanically or thermally sensitive ion channels (Yoo *et al* 2011.) Further experiments are needed to ascertain the temporal window within which performance improvements are obtained. Such experiments should be done by sonicating subjects while they are alert to avoid any confounding effects of anesthesia (Downs *et al* 2015).

Recently, McDannold *et al* (2015) showed that opening the BBB facilitated the blockade of neural activity by GABA in somatosensory cortex of rats. Here, we found that sonication interacted with a low dose of haloperidol, a D2 dopamine antagonist, that was injected 5 minutes prior to behavioral testing. Lower levels of striatal D2 dopamine receptors are associated with reduced motivation and increased impulsivity (Trifilieff & Martinez, 2014.) Previous studies of the effects of haloperidol on response times have reported mixed results depending on species, task and dosage (Brockel and Fowler, 1995, Blokland and Honig, 1999, Kern *et al* 1998). In the current study, low dose haloperidol tended to shorten response time and reduce decision accuracy. Hence, the effects of haloperidol were opposite to those of increasing reward size, consistent with the idea that the behavioral effects of reward may be due to reduced motivation, mediated by striatal D2 dopamine receptors. Haloperidol may inhibit signaling through the indirect basal ganglia pathway, allowing the direct pathway to produce shorter latency movements (Albin *et al* 1989; DeLong *et al* 1990.)

The effects of haloperidol showed an interaction with sonication. This result indicates that FUS can be used in combination with dopaminergic medications to modulate cognitive performance. The results also suggest that the systemic dose of a drug necessary to achieve a desired pharmacological effect may be reduced by increasing BBB permeability through the application of FUS to a targeted brain region, even if the drug in question readily crosses the BBB. This would allow for smaller systemic doses, and thus reduction of potential side effects of currently available drugs for therapies to treat neurological and psychiatric disorders.

There are a few previous studies investigating the effect of FUS without BBB opening on alert subjects performing behavioral tasks. Deffieux and colleagues applied FUS to monkeys performing an antisaccade task by targeting the left frontal eye field (FEF) and the premotor cortex (Deffieux *et al* 2013). Ipsilateral antisaccade latencies were significantly slowed while targeting the FEF but not the premotor cortex. Two other groups investigated the effects of FUS on human subjects (Hameroff *et al* 2013, Legon *et al* 2014). Subjects tested by Legon *et al* exhibited enhanced sensitivity to the frequency of air puffs and improved two-point tactile discrimination while FUS was applied to their somatosensory cortex. FUS was applied to the frontal-temporal cortex in subjects of the Hameroff *et al* study and unlike the other two studies with simultaneous/immediate behavioral testing, results were determined 10 and 40 minutes after application. Subjects reported a significant improvement on the Global Affect test, as well as slightly reduced pain levels 40 minutes after the application of FUS. These studies demonstrate that FUS is capable of affecting the function of the brain depending on the targeting area, while the Hameroff *et al* study shows the effects could be time sensitive. A key difference from the current study is that in the aforementioned studies the BBB remained undisrupted in the targeted region to the knowledge of the experimenters.

Recently, our group applied the FUS BBB opening procedure to awake, behaving monkeys performing a reaching task with variable reward magnitude (Downs *et al* 2015). That study found a slight increase in response time as well as a significant improvement in the accuracy of reaching to visual stimuli during a 2-minute application of FUS and throughout the remaining 2 hours of behavioral testing. McDannold *et al* had previously shown that BBB disruption in the region of the lateral geniculate nucleus did not impair visual acuity (McDannold *et al* 2012). Chu *et al* investigated the effects of BBB opening via FUS opening on somatosensory evoked potentials (SSEPs) and blood-oxygen-level dependent (BOLD) responses when targeting the left primary somatosensory cortex in anesthetized rats (Chu *et al* 2015). Results showed both a decrease in SSEP and BOLD signals within 10 minutes after finishing the FUS procedure with effects lasting up to 7 days. Their results highlighted the impact of sonication parameters utilized, as lower acoustic pressures resulted in little to no neurological effect, while higher acoustic pressures created sustained neurological effects. Our study utilized an acoustic pressure found to be safe during prior studies conducted within our lab, which was greater than the pressure used by Chu and colleagues. The exact mechanisms behind the excitation or inhibition of neurons via FUS is currently unknown, but one theory is that mechanical forces emitted by the transducer during sonication affect mechanoreceptors in the cell membrane (Tyler *et al* 2008, King *et al* 2013, Velling and Shklyaruk, 1988). However, this mechanism is likely to be limited to cases in which the sonication is applied simultaneously during the behavioral testing. Our results, along with the studies conducted by the Hameroff and Chu studies, demonstrate that the effects of FUS sonication can persist after the time of application. Further studies plan to determine the optimal time after FUS application to open the BBB for behavioral modulation. Understanding the relationship between treatment time and behavioral effects will help distinguish the mechanical effect of the sonication from the other potential neurological effects of the BBB being opened at the target region.

## Conclusion

In conclusion, opening the BBB via FUS with microbubbles can have a significant effect on the behavioral responses of monkeys 3-4 hours after the end of the sonication. The BBB opening also facilitated the delivery of a low dose of haloperidol demonstrating that therapeutic doses of a drug can be reduced to mitigate the potential side-effects after opening the BBB at the target region for therapy. Overall, our results demonstrate the potential for FUS to enhance cognitive function.

## Acknowledgements

NIH R01MH059244 (VPF); Kavli Foundation (VPF); Behavior and Brain Research Foundation (VPF); NIH R01AG038961 (EK); NIH R01EB009041 (EK); Wallace H. Coulter Foundation (EK); FUS Foundation (EK); Alfonso Martin Escudero Foundation, Spain (CS). Barrie Foundation, Spain (CS). The authors declare no competing financial interests.

